# Quantifying key parameters of environmental transmission and age-specific susceptibility for *Mycobacterium avium* subspecies *paratuberculosis (MAP)*

**DOI:** 10.1101/2024.11.14.623589

**Authors:** Yuqi Gao, Piter Bijma, Nienke Hartemink, Mart C.M. de Jong

**Affiliations:** Infectious Disease Epidemiology Group, Department of Animal Sciences, Wageningen University and Research, Wageningen, Netherlands; Animal Breeding and Genomics Group, Department of Animal Sciences, Wageningen University and Research, Wageningen, Netherlands

## Abstract

Paratuberculosis is a chronic disease in cows and other ruminants, caused by *Mycobacterium avium subspecies paratuberculosis* (*MAP*). We developed an age-specific dose-response model and two environmental transmission models (Model A and B) to estimate key parameters based on previously published experiments for MAP in dairy cows. In the dose-response model, the age-specific susceptibility decrease rate parameter was estimated at 0.0629◻wk^−1^, suggesting that a previously used parameter of 0.1◻wk^−1^ may have underestimated the infection risks with increasing age. For the transmission models, Model A represents infectivity differences among transiently infectious (*I tr*), low shedding (*Il*), high shedding (*Ih*) individuals by varying transmission rate parameters (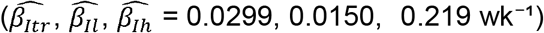) with a standardized constant shedding rate parameter of 2.10◻wk^−1^, whereas Model B captures these differences by varying shedding rate parameters (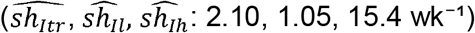) with one transmission rate parameter of 0.0299◻wk^−1^. Although both models have identical best estimates and AIC values, Model B exhibited wider 95% confidence intervals (95% CI). In both models, the MAP decay rate parameter was estimated at 0.150◻wk^−1^, corresponding to a half-life of MAP of approximately 4.61 weeks, which aligns well with previously published values. To better interpret these parameters and understand how different biological assumptions about infectivity influence predicted exposure, we performed scenario analyses examining environmental contamination over time, along with infection probabilities over exposure timing, exposure duration, and the recipient’s age at exposure. All 95% CI are provided in the main text.

## Introduction

Paratuberculosis, caused by *Mycobacterium avium subspecies paratuberculosis* (*MAP*), is a chronic disease affecting cows and other ruminants (1). The disease is known for its long incubation period, imperfect diagnostic tests (especially for early stages), limited vaccination effect, and long survival in the environment, all of which alone or in combination could be the reason why *MAP* is difficult to control and eradicate in cattle(2-7). *MAP* can be transmitted horizontally via contaminated milk or faeces and vertically from dam to offspring (8, 9). A major obstacle in eradicating *MAP* is its environmental transmission, where contaminated environments can cause infections even in the absence of infectious individuals at the location and time of the recipients’ presence.

Understanding environmental transmission is crucial for *MAP* dynamics; therefore, we applied an advanced mathematical model developed by Chang and De Jong (10) to study it. This model, which has proven effective for various diseases (10-13), was not yet used for *MAP*. The model incorporates three key parameters: for, shedding rate, decay rate, and transmission rate. Additionally, if spatial spread is considered, a fourth parameter, for the diffusion rate, is necessary (12). For *MAP* on dairy farms, spatial transmission was simplified by considering three different environments on a dairy farm: the calving house, calf collective pen, and lactating adult pen. This research focuses on estimating the shedding, decay and transmission rate parameters, differentiated for individuals at various infectious states, following the approach of Chang and De Jong (10). The model assumes that infectious animals shed *MAP* into the environment at their shedding rate, and once present, *MAP* contamination level in the environment decays exponentially according to *MAP*’s natural decay rate. The cumulative exposure is determined by integrating environmental contamination level. This approach allows us to capture transmission both in the presence and absence of actively shedding individuals. By estimating all necessary transmission parameters to support the application of this model to *MAP*, our goal is to improve our understanding of *MAP* transmission dynamics and to guide interventions aimed at effectively controlling and potentially eradicating *MAP*.

Accurate assessment of infection risk in *MAP* requires a thorough understanding of age-specific susceptibility. Newborn calves, probably because of their heightened intestinal permeability, are particularly vulnerable to *MAP* during the first 24 hours of life (14, 15). Susceptibility decreases with age, with cows generally considered susceptible only during their first year of life (16). However, infections in adult cows have been observed, but only when exposed to very high levels of *MAP*, such as through oral dosing of 180 mg of cultured MAP or housing with 15 high shedders for 4 years (14, 17). The commonly used exponential model for age-specific susceptibility assumes a decrease rate parameter of 0.1 wk^−1^ (16, 18-20). However, this parameter estimate is a broad estimate and could possibly be improved. Our analysis addressed this gap by developing an age-specific dose-response model and estimating the susceptibility decrease rate parameter using data from fourteen inoculation experiments that include age at exposure. The study aimed to more accurately capture how susceptibility to *MAP* varies with age and dose, and to enhance the precision of infection risk calculation.

This research developed both an age-specific dose-response model and two stochastic environmental transmission models (Model A and B) to estimate parameters and their 95% confidence interval (95%CI) based on fourteen dose-response experiments and two transmission experiments. The parameters include the parameters for susceptibility decrease rate (*g*), median infective dose (*ID*_50_), transmission rate (*β* or *β*_*state*_), shedding rate (*sh*_*state*_ or *sh*), *MAP* decay rate in the environment (*de*), where *state* is either transiently infectious (*Itr*), low shedding (*Il*) or high shedding (*Ih*). This classification of shedding patterns is consistent with the observed shedding patterns from transmission experiments. With the modes and the estimated parameters, we then explored four scenarios: changes in contamination levels over time, timing and duration of exposure, and the recipient’s age at exposure.

## Materials and Methods

### Age-specific susceptibility decrease rate

Literature search was conducted focusing on *MAP* inoculation experiments performed on calves with different age groups. Data were collected from experiments that provided detailed information, including inoculation dose, bacterial strain, number of experimental animals, age at inoculation, inoculation methods, diagnostic methods, and binary results (infected/not infected) (2, 3, 21-35). The initial susceptibility of new-borns is standardized to be 1 and it decreases exponentially with age at a rate of *g. ID*_50_ is defined as the dose at which a newborn calf, when exposed, would have a 50% probability of becoming infected. To estimate *g* and *ID*_50_, we constructed an age-specific dose-response model to calculate relative susceptibility (*rS*(*age*)), based on exponential function (Equation 1). The maximum likelihood method is applied on a stochastic model with the number of new infections following a binomial distribution, with *S* trials and a dose/age dependent infection probability *p*(*dose*,*age*) (Equation 2). In the model, *g* is regarded as an adjustment factor influencing *ID*_50_, which implies that older individuals require a higher dose of *MAP* to attain a 50% chance of infection compared to younger individuals.

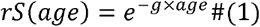

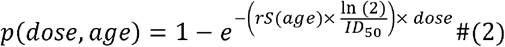

Besides, to calculate the infection probability over an exposure period, where the age increases simultaneously over time, the average relative susceptibility can be calculated as 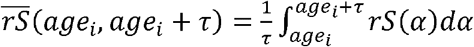. This calculation provides the mean susceptibility over the time interval (*i, i*+ τ), accounting for how susceptibility changes as the individual ages from *age* _*i*_ to *age i* +*τ*.

### Transmission rate parameters

*MAP* experimental transmission data were obtained from experiments conducted by van Roermund et al. (36), which involved 12 cows and 20 calves, and aimed to replicate real-world cow-to-calf and calf-to-calf transmission scenarios. In the initial phase of the experiment, six inoculated adult cows were placed in a row, and five 1-week-old naive calves(*S*) were placed in between two successive cows. The positions of calves were altered randomly every two weeks over the period of the three-month experiment, ensuring each calf was equally exposed to all pairs of cows in the row. Sampling of all individuals occurred once every two weeks, using three diagnostic methods, i.e., serum ELISA, bacterial culture, and interferon-gamma tests. Cows with at least one positive result (based on faecal culture) of more than 100 colonies of *MAP* per tube (0.16 g faeces) out of the seven tests were categorized as high shedders (*Ih*), while those with at least one positive result or no positive result were categorized as low shedders (*Il*) or non-infectious, respectively. This classification was also applied throughout our modelling.

Subsequently, in the second phase of the experiment, the previously exposed five calves (including susceptible, latently infected (*L*), and transiently infectious calves (*Itr*)) were brought into contact with five new 1-week-old naive calves for a period of three months. Within the calf pen, all 10 calves had unrestricted contact with each other. Like before, the three diagnostic methods were conducted once every two weeks on each calf. The whole experiment was repeated once, with again 6 cows, 5 naïve calves, and these were again brought in contact with 5 additional calves, to verify consistency and accuracy of the findings. Regular pen cleaning was performed weekly, without additional disinfection. All calves were fed artificial milk replacer, eliminating the possibility of transmission through contaminated milk. A summary of the original data from published experiments is provided in S1 File.

Based on data from these published experiments, we constructed two stochastic environmental transmission models (A and B) to estimate three key parameters in *MAP* transmission dynamics: i.e. those for shedding rate, transmission rate, and *MAP* decay rate. According to the model assumptions, the environmental contamination level (*Estate*(*t*)) is determined by the presence and shedding rate of infectious individuals at different states (*state* = *Itr, Il* or *Ih*), as well as the pathogen’s decay rate in the environment (Equation 3). Specifically, Model◻A captures infectivity differences through varying transmission rate parameters, while Model ◻ B captures them through varying shedding rate parameters. The detailed schemes for both models are provided in S2 ◻ File.

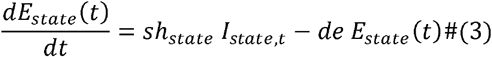

with *E*_*state*_(0) =0

By solving Equation 3, *E state* (*t*+ *τ*) at the end of the interval (*t*,*t* + *τ*) can be expressed explicitly, when *E*_*state*_ (*t*) and *I*_*state,t*_ are known :

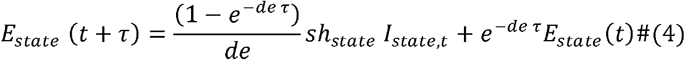

In the environment, the total contamination is contributed by all infectious individuals in the environment. As we assume individuals’ infection states remain constant over a time interval (*i, i*+*τ*), the total force of infection λ*tot*_*i,i+*τ**_ is calculated by summing contributions from each individual *m* in the infectious compartment *I*_*i*_, where *m*∈ {1,2…1*i*}. Specifically, an individual *m* contributes 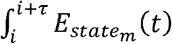 to the environment, which reflects the amount of MAP shed during the interval (*i, i*+*τ*), when the instantaneous contamination level 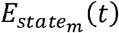 is given by Equation 4. For the force of infection rate calculation, in the transmission-rate-based Model A, this quantity (of accumulated contamination) is multiplied by the corresponding transmission rate parameter 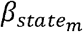, representing the infectivity level of that shedding state. Alternatively, in the shedding-rate-based Model B, shedding parameters vary by *state*_*m*_ and a constant transmission rate applies to all states.

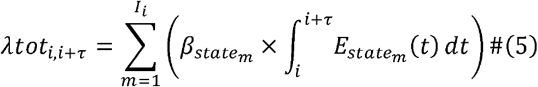

Then, the infection probability (*p*_*i,i+*τ**_) for a recipient at age (*age*_*i*_, *age*_*i*_ +*τ*), during time interval (*i, i*+ *τ*) equals:

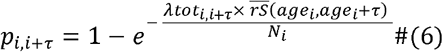

Here, *N*_*i*_ denotes the size of environment (37).

Although, each individual was tested every two weeks, we produced weekly model outputs (*τ* ◻= ◻1◻wk) to capture new infections while the underlying model runs in continuous time. We defined the time of infection as the midpoint between the last negative test and the first positive test. However, in experiments B1 and A2, six calves remained negative throughout the contact period but tested positive only during the subsequent individual housing period. In these cases, we assumed the calves bypassed the transient shedding stage and entered a latent phase directly. With no further data available, we assumed the midpoint of the experiment as the most likely time of infection for these animals. The maximum likelihood method is used to fit the model to observed infection events, with the estimates determined at the minimal Akaike Information Criterion (AIC) value. The 95%CI are determined by adding 2 to the minimum AIC value(38). Equations for maximum likelihood estimate 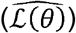 and AIC are listed below:

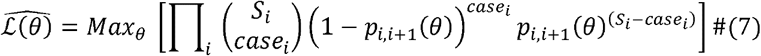

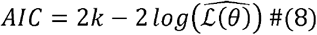

with θ= {*de*,*β*_*Itr*_, *β Il*, *β Ih*} in model A, the shedding rate *sh* is a function of *de* as follows: 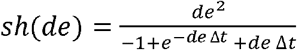, with Δ*t*=1 wk (10). Or, in model B, with *θ*= {*de*,*β*_*Itr*_,*β Il*, *β Ih*} and 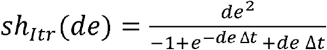, with Δ*t*=1 wk. The number of unknow parameters *k*, equals 4.

As shown above, *β Itr* in model A equals to *β* in model B, and *sh* in model A equals to *sh*_*Itr*_ in model B. This is because *Itr* is the reference infectious state in both models, which can be explained by the unit definition of the model. The unit of *E*_*state*_(*t*) differs in scale from the dose unit (e.g. CFU for *MAP*). Instead, it is standardized such that, in a clean environment, the *Exposure*, from one infectious individual at state *Itr* over a week is set to one unit 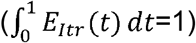. This standardization ensures a consistent interpretation of the transmission rate parameter and shedding rate parameter across this research and other studies with different transmission mechanism assumptions(10).

### Model fit evaluation

Model fit is assessed by comparing the predicted infection probabilities with the actual observations from the experiments. The experiment begins at time 0, and at any subsequent time point *t, p*_0,*t*_ can be calculated using the Equation 6. The total infection probability over the whole experiment duration and the expected number of new infections can be calculated (*E*(*Newcase*) = *S*_0_ × *p*_0,*end of experiment*_). This evaluation helps to determine how accurately the model reflects the true dynamics of infection transmission and provide insights into how the underlying infection process may look like in the real-world experiments.

### Scenario study

In this research, a scenario study was included to show the interpretation of model A and B, and the estimated parameters under more real-world conditions. The study focuses on four key drivers of transmission: (1) Duration of contamination: examining the impact of retaining an infectious individual (at state *Itr, Il* or *Ih*) in a clean environment, simulating changes in the contamination level over time (*E*_*state*_(*t*)), the force of infection rate over time(*λ*_*state*_(*t*)) and calculating time (*t*_*eq*_) required to reach equilibrium phase. (2) Timing of exposure: calculating infection probabilities associated with short-term exposures before equilibrium (when *E*_*state*_(*t*) is still increasing) and after equilibrium (when *E*_*state*_(*t*) has stabilized). To ensure meaningful comparisons, the exposure interval is set to 1 week, with the susceptible individual’s age at 0 weeks at the start of exposure. (3) Duration of exposure: Investigating how varying exposure durations influence infection probability over time. Similarly to Scenario 2, the assumptions are that *E*_*state*_(*t*) is stabilized and that the recipient is fully susceptible at the start. (4) Age of the recipient: Predicting how changes in the recipient’s age, along with corresponding variations in susceptibility, affect infection probability, assuming *E*_*state*_(*t*) is stabilized and the exposure duration is fixed at 1 week. For both Scenario 3 & 4, the total simulated duration was based on the maximum susceptible age of 52 weeks. Table 1 provides a summary of the scenario setups, variables, variation ranges, and corresponding equations.

**Table 1.** Overview of Scenario Study Parameters and Objectives.

| Scenario | Infectious state | Exposure starts time (Week) | Length of exposure interval (Week) | Recipient's age at start of exposure (Week) | Output |
| --- | --- | --- | --- | --- | --- |
| 1 | $I_{tr}, I_l$ or $I_h$ | $t, t \in [0, 52]$ | - | - | $E_{state}(t)$ and $\lambda_{state}(t)$ |
| 2 | $I_{tr}, I_l$ or $I_h$ | $x, x \in [0, 52]$ | 1 | 0 | $p_{x,x+1} = 1 - e^{-\beta_{state} \times \int_x^{x+1} E_{state}(t) dt \times \overline{rS}(0,1)}$ |
| 3 | $I_{tr}, I_l$ or $I_h$ | $eq^*$ | $y, y \in [0, 52]$ | 0 | $p_{eq,eq+y} = 1 - e^{-\beta_{state} \times \int_{eq}^{eq+y} E_{state}(t) dt \times \overline{rS}(0,y)}$ |
| 4 | $I_{tr}, I_l$ or $I_h$ | $eq^*$ | 1 | $z, z \in [0, 52]$ | $p_{eq,eq+1} = 1 - e^{-\beta_{state} \times \int_{eq}^{eq+1} E_{state}(t) dt \times \overline{rS}(z,z+1)}$ |

\* $eq$ can be set to any time point after $E_{state}(t)$ reaches its equilibrium ( $t_{eq}$ ), which will be informed by the outcome of scenario 1.

All calculations and simulations were performed using Mathematica 14.0, with machine precision applied throughout the computations to ensure numerical accuracy. All data and code used for parameter estimation, simulation, model fitness analysis and scenario analysis are available in a Git repository at https://git.wur.nl/yuqi.gao/map_para_esti.

## Results

### Age-specific susceptibility decrease rate

The dose-response model was used to estimate infectious Dose 50 (*ID*_50_) and age-specific decrease rate (*g*) parameters from *MAP* inoculation experiments in calves of different ages. The fit of the non-linear model, based on an exponential function, results in an *ID*_50_ estimate and its 95%CI of 2.75 × 10^6^ (1.42 × 10^6^, 4.08 × 10^6^) CFU, and a *a* estimate and its 95%CI of 0.0629 (0.0446, 0.0811) wk^−1^. We plotted the predicted age-specific dose-response plot, overlaid with the original observations (dot sizes proportional to sample sizes) (Fig. 1).

**Fig 1.**
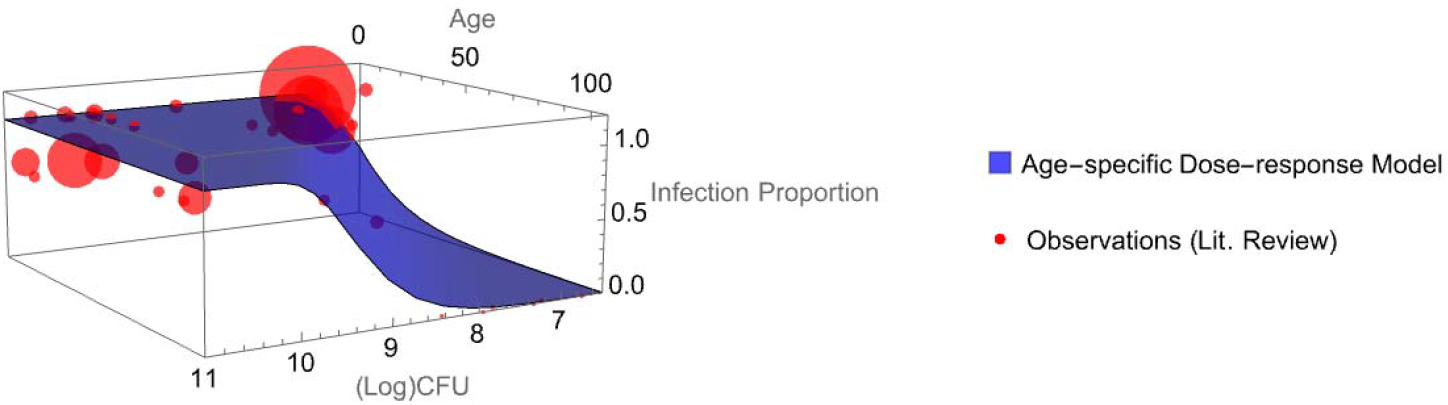
Age-Specific Dose-Response Model Prediction. This figure illustrates the predicted age-specific dose-response relationship (blue surface), overlaid with observed data points scaled by sample size (larger dots = larger recipients’ group).

For model fit evaluation, the 95% prediction interval (PI)—accounting for both model uncertainty and the binary nature of infection—encompassed all observed data. Additionally, we achieved the coefficient of determination (R^2^) of 0.797 and adjusted R^2^◻of◻0.796, which means our model explains nearly 80% of the variance. The minimal gap between R^2^ and adjusted R^2^ further suggests minimal overfitting. Relatively susceptibility (*rS*(*age*)) decreases exponentially with age at a rate of 0.0629 wk^−1^ and the average susceptibility 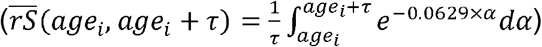 within age range (*age*_*i*_,*age*_*i*_ + *τ*) is quantified by integrating the continuous *rS*(*age*) function and diving it by the interval length.

As shown in Fig. 2, our model estimates a slower decrease rate in susceptibility with age compared to the previously published rate of 0.1 wk^−1^. This suggests that susceptibility decrease rate with age has been overestimated, while the infection risk has been underestimated.

**Fig 2.**
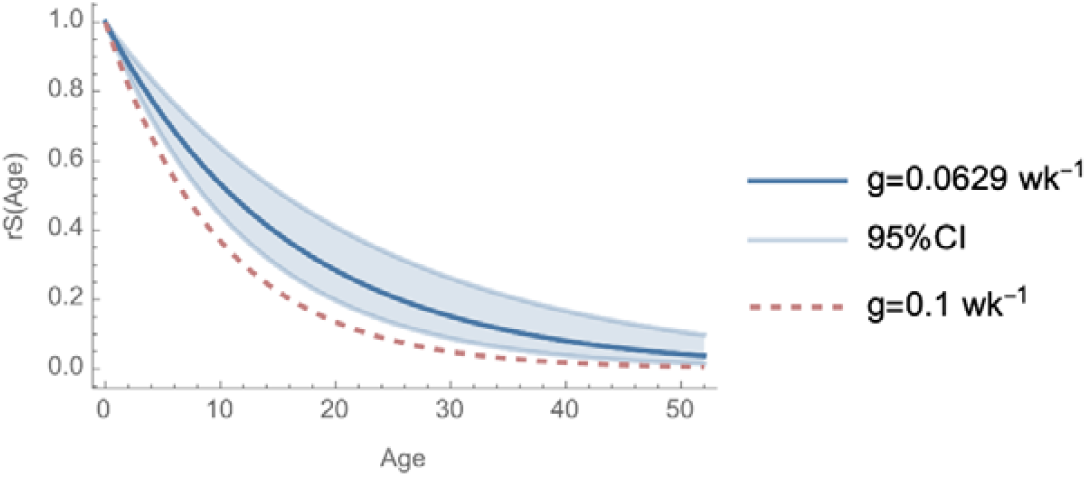
Comparison of Susceptibility Decrease with Age. The solid line and shaded region represent the relative susceptibility decrease (*rS*(*age*)) with age based on our model-estimated rate (*g* = 0.0629 wk^−1^) and its 95% confidence interval. The dashed line shows the relative susceptibility decrease using the previously published rate of 0.1 wk^−1^.

### Transmission parameters estimation

The environmental transmission model parameters were estimated from experimental cow-to-calf and calf-to-calf transmission data using maximum likelihood estimation. These included the decay rate parameter estimate for *MAP* in the environment 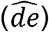 transmission rate parameter estimates, which were state-specific 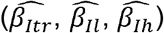 in Model A, and constant (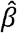) in Model B; and shedding rate estimates, which were constant 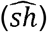 in Model A, and state-specific 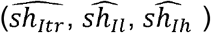 in Model B. In both models, the reference infectious state is *Itr*, which means that 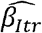 model A is equivalent to 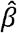 in model B, and 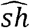 in model A is equivalent to 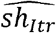 in model B. Profile likelihood visualizations for each parameter are shown in Fig 3, with detailed results in Table 2.

**Table 2.**
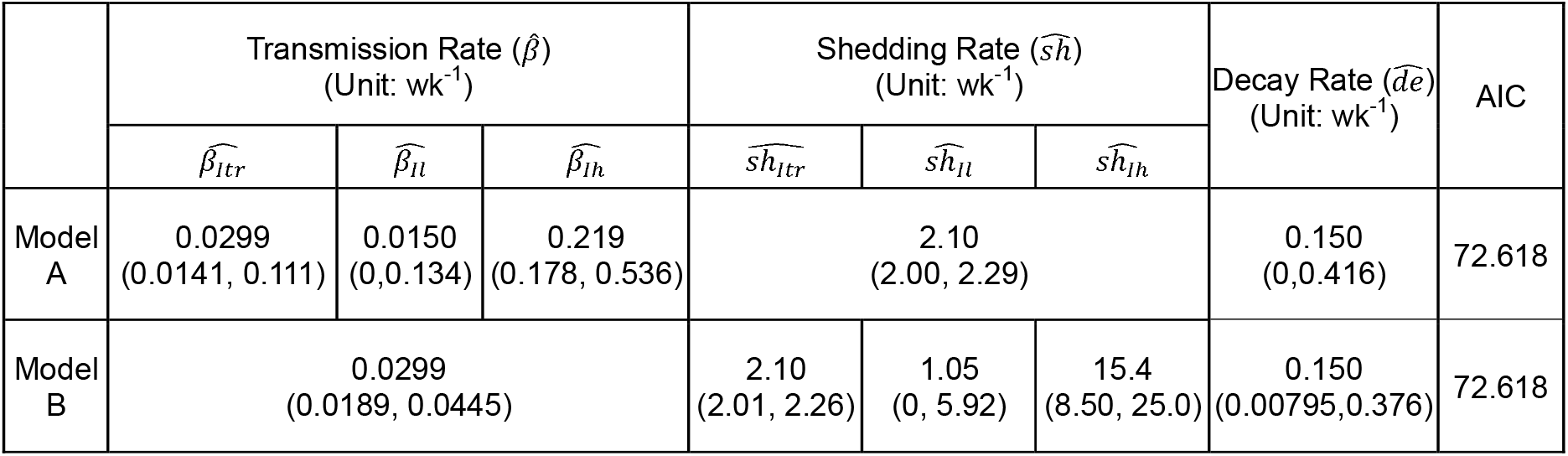
Profile Likelihood Estimates and 95% Confidence Intervals for Transmission, Shedding, and Decay Rate Parameters in Model A and B.

**Fig 3.**
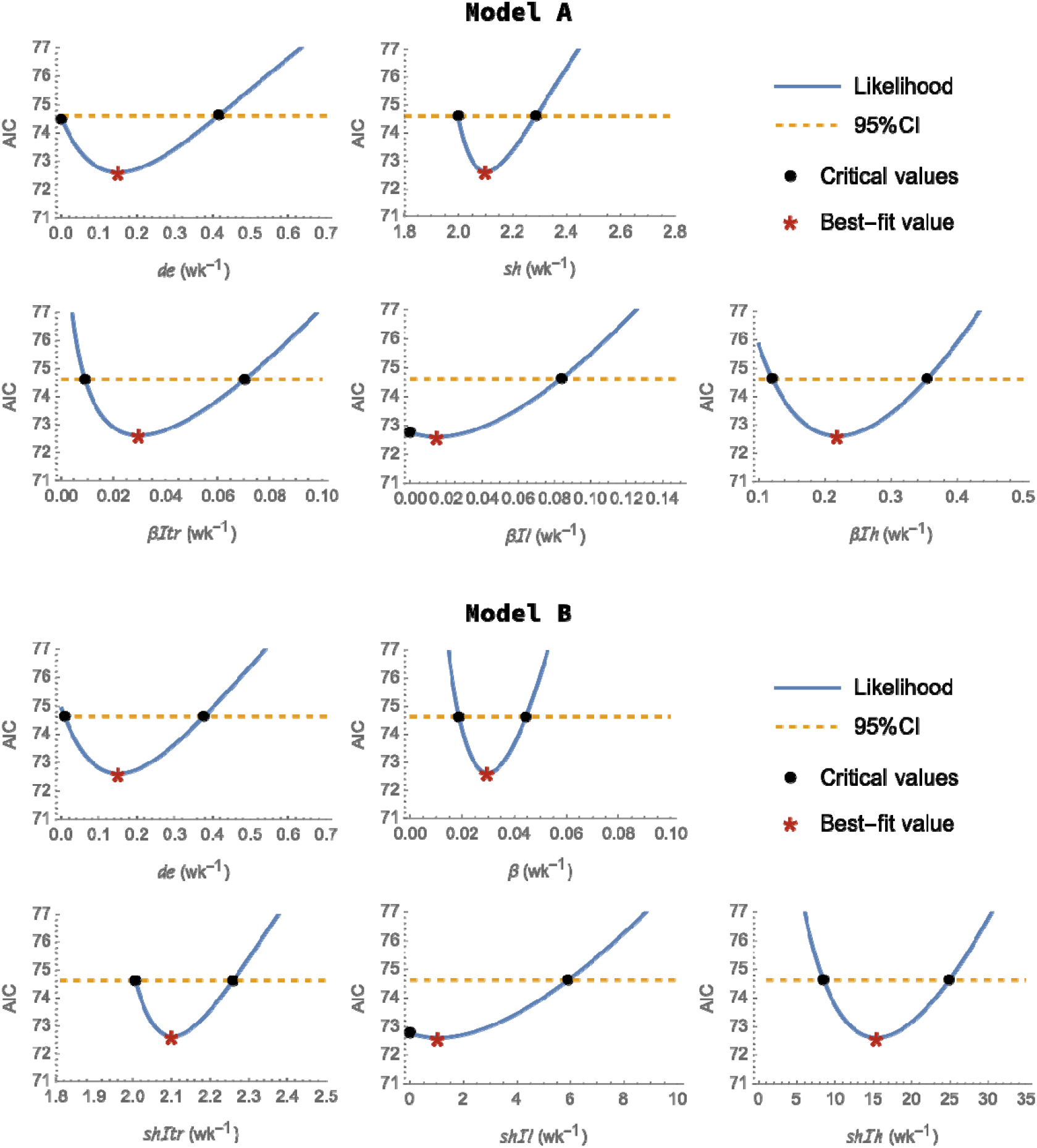
Profile-Likelihood Plots for Shedding, Transmission, and Decay Rate Parameters. The upper panel represents results from Model A, where the lower panel represent Model B. The best estimate for each parameter corresponds to the lowest AIC value, marked by a red star. The black dots represent the critical values for the 95% confidence interval, which is defined by minimum AIC+2 (the dashed line).

### Model fit evaluation results

Model fit is assessed by comparing predicted infection probabilities with actual experimental observations. During Experiment A1, 3 high shedders and 3 low shedders were observed; during Experiment A2, 4 transient shedders were present. During Experiment B1, 1 high shedder, 3 low shedders, and 2 non-infected cows were observed, while during Experiment B2, no infectious individuals were present. As detailed in the Materials & Methods section, in Experiments A1 and B1, 6 cows were housed in individual pens, with 5 recipient calves placed between each pair of cows, and calves’ positions were randomly switched every two weeks. For simplicity, this setup was treated as a “pairwise” experiment, where one cow and one calf are paired. As there was no specific information regarding the location of cows and calves in the experiment A1 or B1, the infectivity of the shedder in each pair was considered as the average across all 6 infected cows. In experiments A2 and B2, five shedders and five recipient calves were placed in the same pen, allowing for unrestricted contact between all animals. Based on the number and infectious state of individuals in each experiment, the environmental contamination levels (*E*_*state*_(*t*)), population force of infection (*λtot*_0,*t*_), and infection probabilities (*p*_0,*t*_) were calculated according to Equations 3-6.

As shown in Fig 4, the curve and shaded area (left axis) represent the predicted infection probabilities over time (0, t), while the bar plot (right axis) displays the observed data from the experiments. The visualization indicates a good fit of the model, as the predictions aligns with the observed data, demonstrating that the model effectively captures the dynamics of infection transmission. Besides, the total infection probabilities over the whole experiment duration and the predicted number of infections were calculated. In both model A and model B, to three decimal places, the predicted overall infection probabilities for experiment A1, A2, B1, B2 are 97.8%, 45.2%, 76.3%, 0.00%. As there are five recipients in each experiment, the expected number of new infections is calculated to be 4.89, 3.81, 2.26, 0, respectively, while 5, 4, 2, 0 infections were observed.

**Fig 4.**
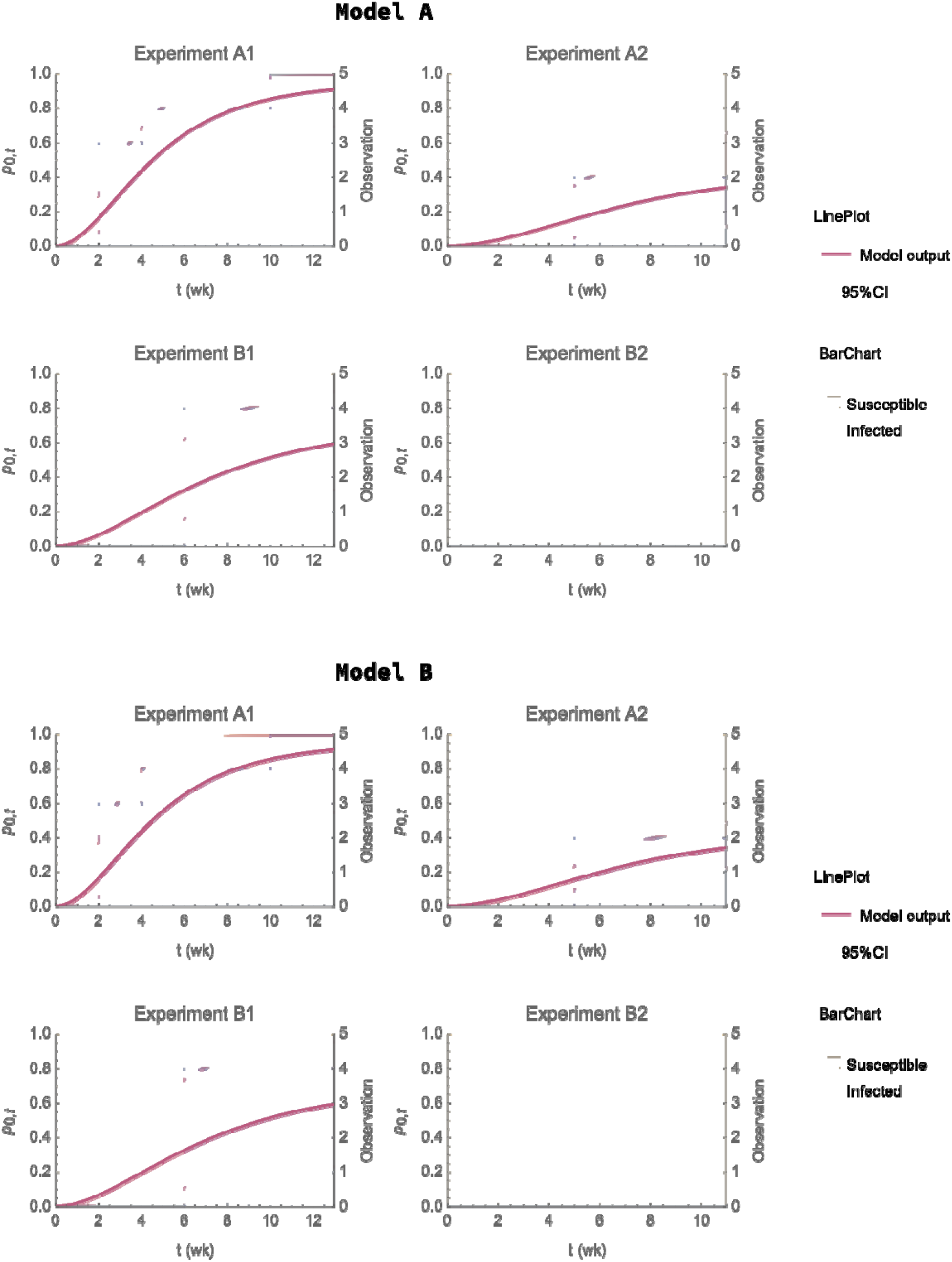
Comparison of Predicted Accumulative Infection Probabilities with Experimental Observations in Model A and B. The upper panel represents results from Model A, where the lower panel represent Model B. The four plots in each panel show parallel cow-to-calf experiments (A1 & B1) and parallel calf-to-calf experiments (A2 & B2). The solid curve and shaded area represent the predicted accumulative infection probabilities and their 95% confidence interval from the start of experiments(t=0) to time t. The blue bars indicate the number of newly infected individuals observed in the experiments. The x-axis denotes time in weeks, with the left y-axis showing infection probability and the right y-axis showing the number of susceptible and infected individuals.

### Scenario study results

To better interpret these parameters and explore how different biological assumptions about infectivity influence predicted exposure, we conducted a scenario study. Specifically, we simulated changes in environmental contamination level and force of infection rate over time and evaluated the effects of exposure timing, duration, and the recipient’s age on infection probability (Table 2.). However, the decay rate exhibited an extremely lower bound (0 for Model A and 0.00795 for Model B), making it difficult to discern the impact of the other parameters. To focus on the comparison of Model A and B, we therefore fixed the decay rate at its best-estimate value. Consequently, the 95%CI present later for scenario analysis results reflect only the uncertainty from the shedding, transmission, age-specific susceptibility decrease rate parameters. Figures 5 and 6 present the results of this scenario study.

**Fig 5.**
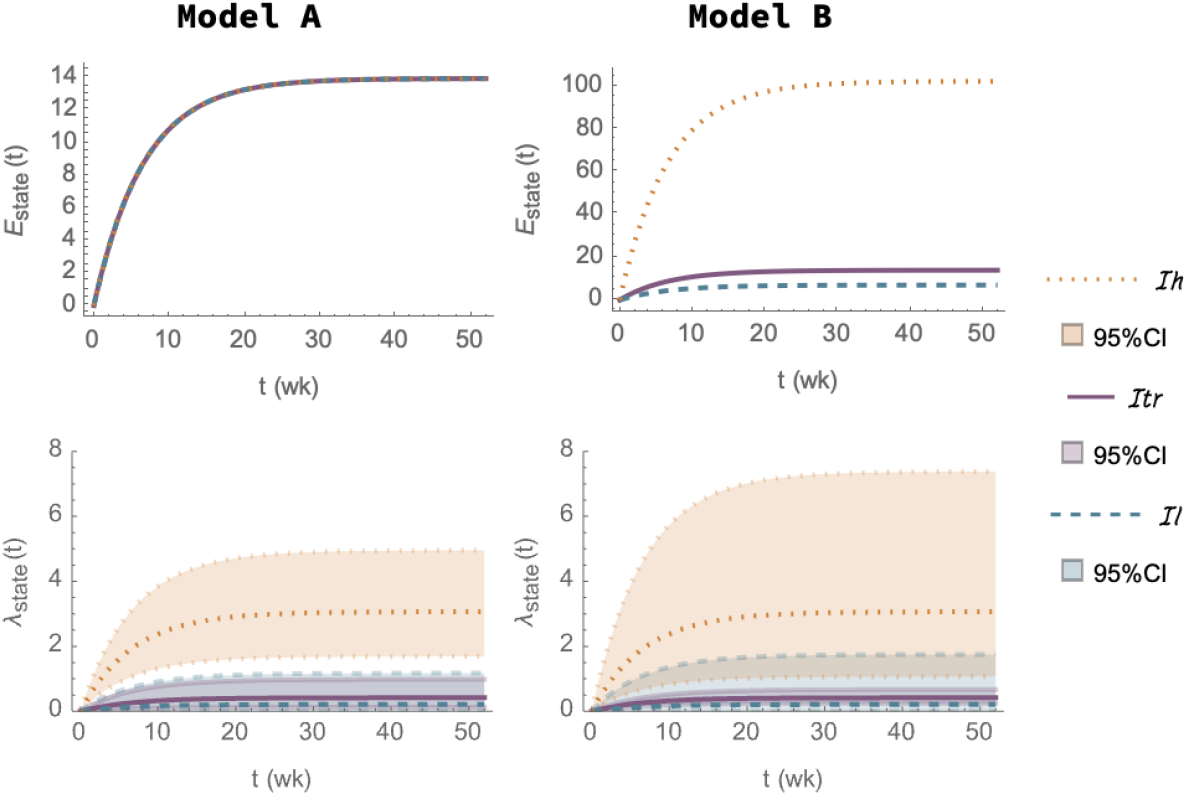
Result of Scenario 1: Changes in *E*_*state*_(*t*) and *λ*_*state*_(*t*) over time while there is one Infectious individual in states *Itr*, It or *Ih*: In Model A (left column), infectious individuals in different states have the same shedding rate, while in Model B (right), *E*_*state*_(*t*) increases at different rates due to varying shedding rates among infectious states. The natural decay rate of MAP is constant in both models.

**Fig 6.**
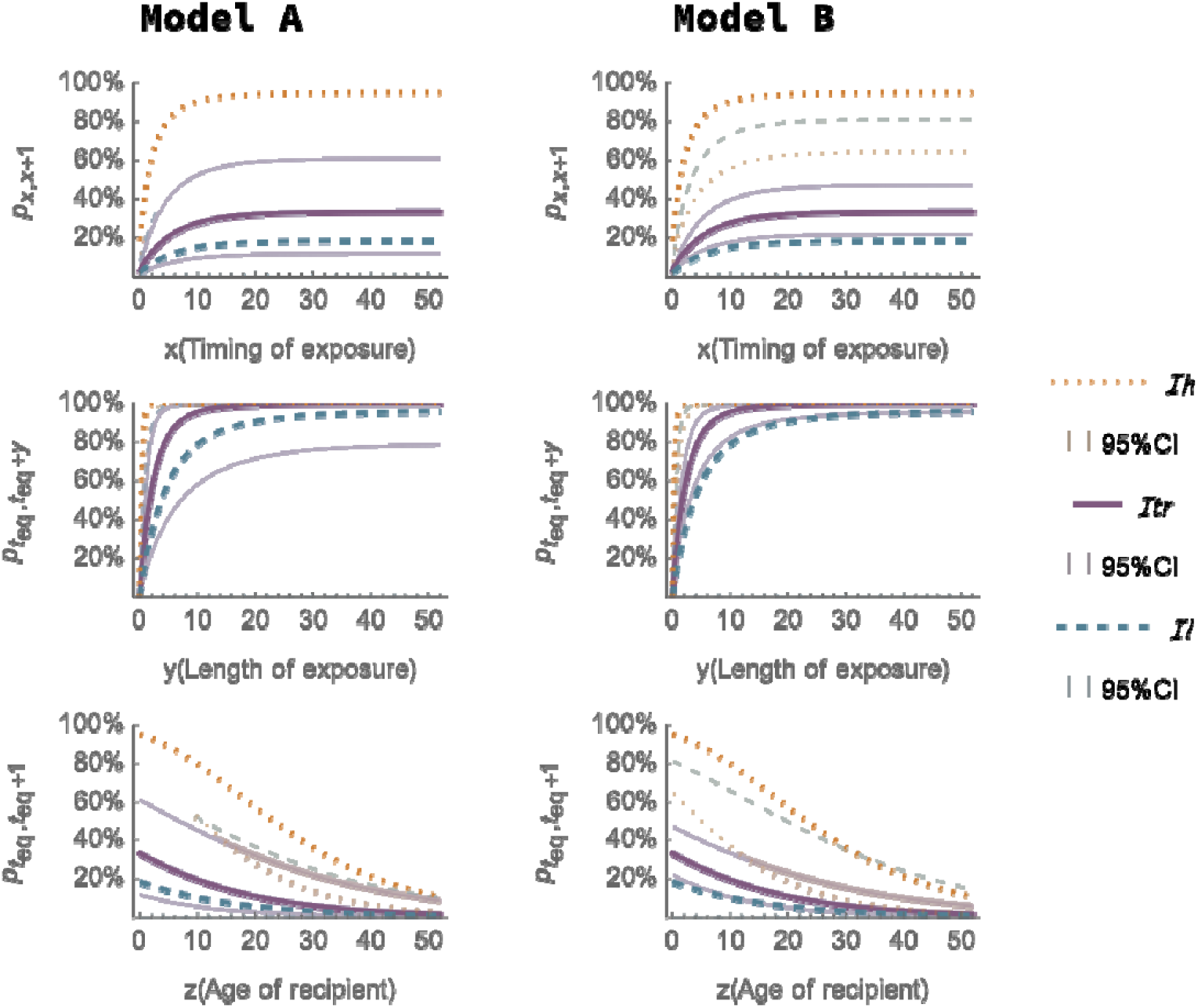
Results of Scenarios 2,3,4: Upper: Predicted infection probabilities for a fully susceptible individual exposed to the contaminated environment at time (for [0,52]) after the start of contamination (=0), with each exposure lasting for a 1-week interval; Middle: Predicted infection probabilities for a fully susceptible individual exposed to the contaminated environment at equilibrium, across varying exposure durations *y* (for *y* ∈ [0,52]); Lower: Predicted infection probabilities for susceptible individuals of age *z* (for *z* ∈ [0,52]) exposed to the contaminated environment at equilibrium, over a 1-week exposure interval. Solid, dashed, and dotted lines represent the infectious individual at states *Itr, Il*, and *Ih*, respectively. The shaded areas represent the corresponding 95% confidence intervals, calculated based on the 95% confidence intervals of transmission rate parameters, shedding rate parameter and the age-specific susceptibility decrease rate.

In Scenario 1, *E*_*state*_(*t*), λ_*state*_(*t*), and *t*_*eq*_ were calculated for environments with the presence of an infectious individual in states *Itr, Il* or *Ih*. The 95%CI were derived from uncertainties in transmission and shedding rate parameters. As shown in Fig 5, *E*_*state*_(*t*) rapidly increase and then stabilize at equilibrium, requiring 30.6◻weeks toreach 99% of the environmental contamination equilibrium. Following the methods in Materials and Methods, we set *teq* =60◻wk for subsequent analyses.

Scenario 2 assessed the infection probability for exposing a fully susceptible individual to the previously defined three types of environments for 1 week, with the variable being the timing of exposure *x*. As shown in Fig 6-upper row, the infection probabilities firstly increased and then stabilized at 33.2% for *Itr*, 18.3% for *Il*, and 94.9% for *Ih*. In Scenario 3, extending the exposure time *y* results in a rapid increase in infection probabilities. For *Ih*, the infection probability approaches 100% after approximately 1.83 weeks. However, for *Itr* and *Il*, the probabilities do not reach 100% due to the decreasing susceptibility over age/time but still reach 99.8% and 95.9% respectively by the end of 52 weeks. In Scenario 4, infection probabilities decrease with age . By the end of calfhood (52 weeks old), after one week of exposure to a stabilized contaminated environment, the predicted infection probabilities are: 1.53% for, 0.767% for, 10.7% for . Therefore, we suggest heifer/cow above one year should also be considered susceptible. In all four scenario analysis, although model A and B produce the same mean prediction values, Model B has wider 95%CI for low and high shedders but a smaller 95%CI for transient shedder.

This research aims to improve understanding of *MAP* transmission dynamics through a novel mathematical model, which has not been previously applied to *MAP*. While comparisons with earlier studies are insightful, differences in transmission mechanism assumptions do complicate direct comparisons. However, the force of infection rates and infection probabilities can still be helpful for evaluating our findings alongside previous research. Therefore, we used a different and commonly used dose-response transmission model (19) and the parameters relevant to that model to replicate the model fit evaluation and scenario analyse. Both models fit the cow-to-calf experimental observations reasonably well; but a significant difference is noticed in calf-to-calf transmission. The predicted number of new cases over the entire period of experiments A1, A2, B1, and B2, based on dose-response model were 5.00, 0.00, 3.98, and 0.00, respectively. These predictions differ from the observed values (5, 2, 4, and 0, respectively), suggesting that the dose-response transmission model may not adequately capture calf-to-calf transmission. Detailed model descriptions, results, and discussions are provided in S4 File.

## Discussion

The aims of this research were: firstly, to develop an age-specific dose-response model to study changes in recipient susceptibility over age; secondly, to construct an environmental transmission model to quantify the parameters for the transmission dynamics of *MAP*, i.e. estimate the shedding rate parameter, transmission rate parameter and *MAP* decay rate parameter. A scenario study followed, focusing on four objectives: changes in contamination levels over time, the timing of exposure, the duration of exposure, and the recipient’s age. Maximum likelihood estimation was applied to achieve the best estimates, and model fit was assessed. Additional details, including a summary of original exposure data from published experiments, culture-based data on *MAP* viability in cattle excretions, and comparisons with transmission parameters from previous research, are provided in supplementary materials.

Research has suggested that calf-to-calf transmission may be underestimated (39-42). In this study, we identified that this underestimation could be partly due to the use of a rounded value of 0.1 for the age-specific susceptibility decrease rate. Based on the exposure data of fourteen experiments, we estimated a decrease rate parameter of 0.0629 wk^−1^ (95% CI: 0.0446, 0.0811 wk^−1^). As shown in Fig. 2, starting from a standardized average susceptibility of 1 for newborns, this value declines to 0.5 after 11 weeks and further drops to 0.0380 by 52 weeks of age. In contrast, the commonly used rate of 0.1 wk^−1^ results in susceptibility dropping to 0.5 at 6.93 weeks and to 0.00552 by 52 weeks. This suggests that if a 52-week-old calf is exposed to a stabilized contaminated environment from *Ih* for one week, the expected infection probability would be 0.927%, which is significantly lower than our estimate of 10.7%. Our prediction aligns more closely with observations of new infections in heifers and older calves (2, 22), and could therefore improve the evaluation of calf-to-calf transmission and inform more effective interventions. Additionally, to verify the estimated 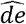, we compare it with culture-based data on *MAP* viability in cattle excretions. For instance, in faecal pellet samples collected from boxes in the 100% shade treatment at 10-22°C, the decay rate was estimated to be 0.189 wk^−1^. Similarly, in cattle feces at site Armidale, Bathurst, Condo Bolin, under temperatures of 3.6–19.7°C, 6.8–19.8°C, 10.2–24.4°C, the decay rates were 0.118 wk^−1^, 0.242 wk^−1^, and 0.242 wk^−1^, respectively. These values provide a range for comparison with the model’s predictions(7, 14, 43-49). Further details can be found in S3 File.

Transmission parameters were estimated using data from cow-to-calf and calf-to-calf transmission experiments. The study applied two models—Model A and B—to capture variations across different infectious states (*Itr, Il, Ih*) in *MAP* transmission. Although Models ◻ A and ◻ B yield identical point estimates and AIC values, Model ◻ B consistently shows wider 95% confidence intervals. One likely explanation is the difference in parameter scaling. In Model ◻ A, infectivity is captured through transmission rates with a standardized shedding rate, whereas in Model ◻ B, the shedding rates vary while transmission remains constant. Because shedding rates in Model ◻ B can span a broader numerical range, small relative changes in shedding can produce large changes in the predicted outcomes, leading to wider confidence intervals.

To interpret the estimates, a scenario study was conducted. The results indicate that *MAP* accumulates quickly in the environment and takes approximately 30.6 weeks to reach 99% of its equilibrium value. At equilibrium, the highest *E*_*state*_ (*t*) is associated with *Ih*, followed by *Itr* and *Il*, though the difference between the latter two is much smaller. However, *Il* exhibits a larger 95% CI, and more importantly, *Il* could pose a greater overall hazard due to its longer shedding period (50). Besides, the scenario 3 shows that the exposure duration greatly affects infection probability, assuming exposing a fully susceptible recipient in a stabilized contaminated environment, the infection probabilities reach 100% for *Ih* after 3.31 weeks, 95.9% for *Il* after 52 weeks and 99.8% for /tr after 52 weeks. The probabilities for *Itr* and *Il* didn’t reach 1 because increasing exposure duration also means the recipient’s age increases, leading to a decrease in relative susceptibility.

In our study, the transmission rate in model A (or shedding rate in model B) for transient shedders is roughly twice that of low shedders; however, experimental data suggest only about half as many infected animals enter the transient shedding phase compared to the low-shedding phase. For example, in Experiments ◻ A1 and ◻ B1, nine calves were infected, but only four began shedding as *Itr* in Experiment ◻ A2. Among all 11 infected calves (nine from A1 & B1 and two from A2 & B2), nine eventually began shedding during the three-year individual housing period after transmission experiments. These observations align with the transient transmission period commonly assumed in modelling studies, where animals briefly shed before entering latency and later resuming low or high shedding. However, when infection histories are reconstructed solely from diagnostic results, distinguishing transient shedders from low shedders—especially in animals infected later in life—can be challenging. We therefore recommend that future backward inference models carefully account for this concern.

A limitation of the current study is the absence of a dedicated environmental transmission experiment—specifically, one that introduces susceptible individuals into a contaminated environment without the presence of infectious individuals. Conducting such an experiment in the future could improve the accuracy of parameter estimation. Besides, the application of the estimated parameters in real farm models is more complex than in controlled lab experiments. On farms, a larger number of infected and susceptible animals at varying states and ages are involved, alongside continuous contamination processes, limited diagnostic capabilities, and more challenging hygiene management due to larger spaces. Next, multiple interventions are widely applied, further complicating quantifying the transmission dynamics. Future research should develop a continuous-time, age-structured compartmental stochastic model that simulates *MAP* transmission at the farm level. Although several simulation models already exist, the refined transmission parameter estimates from this research have the potential to enhance simulation accuracy and improve overall reliability. Such a model would incorporate factors like age cohorts, animal grouping and movement, farm management, environmental contamination, and real-world interventions.

## Supporting information

Supplemental materials S1-S4 files, and will be used for the link to the file on the preprint site

