## Supplemental materials S1-S4 files, and will be used for the link to the file on the preprint site for "Quantifying key parameters of environmental transmission and age-specific susceptibility for *Mycobacterium avium* subspecies *paratuberculosis (MAP)*": S1 File.docx

**S1 File. Collected experimental data from previous research.**

The exact time of infection is unknown, but it can be inferred from diagnostic results. In this research, we assume that infection occurs between the last negative and the first positive fecal culture result. For instance, if calf 5628 tested negative in week 9 and positive in week 11, the infection time is assumed to be at week 10. In Experiment A1, infected calves tested positive during the exposure period, and detailed information was provided to infer the infection process. However, in Experiments A2 and B1, only the total number of infected susceptible individuals was known. These individuals tested negative during the exposure period but were found to be positive later in their respective individual houses. For the latter two cases, we assumed that the individuals became infected during the middle of the experiment. In experiment B2, no infectious individuals were present, and no new infections were observed. Therefore, it was excluded from the parameter estimation.

At the end of period 1, any individuals that remained susceptible could still become infected during period 2. So, both recipients carried over from period 1 and the newly introduced recipients could become infected. However, their ages and levels of susceptibility differed.

**Table S1. Summary of Observations and Inferred Time of Infection from Original Experimental Data (1)**

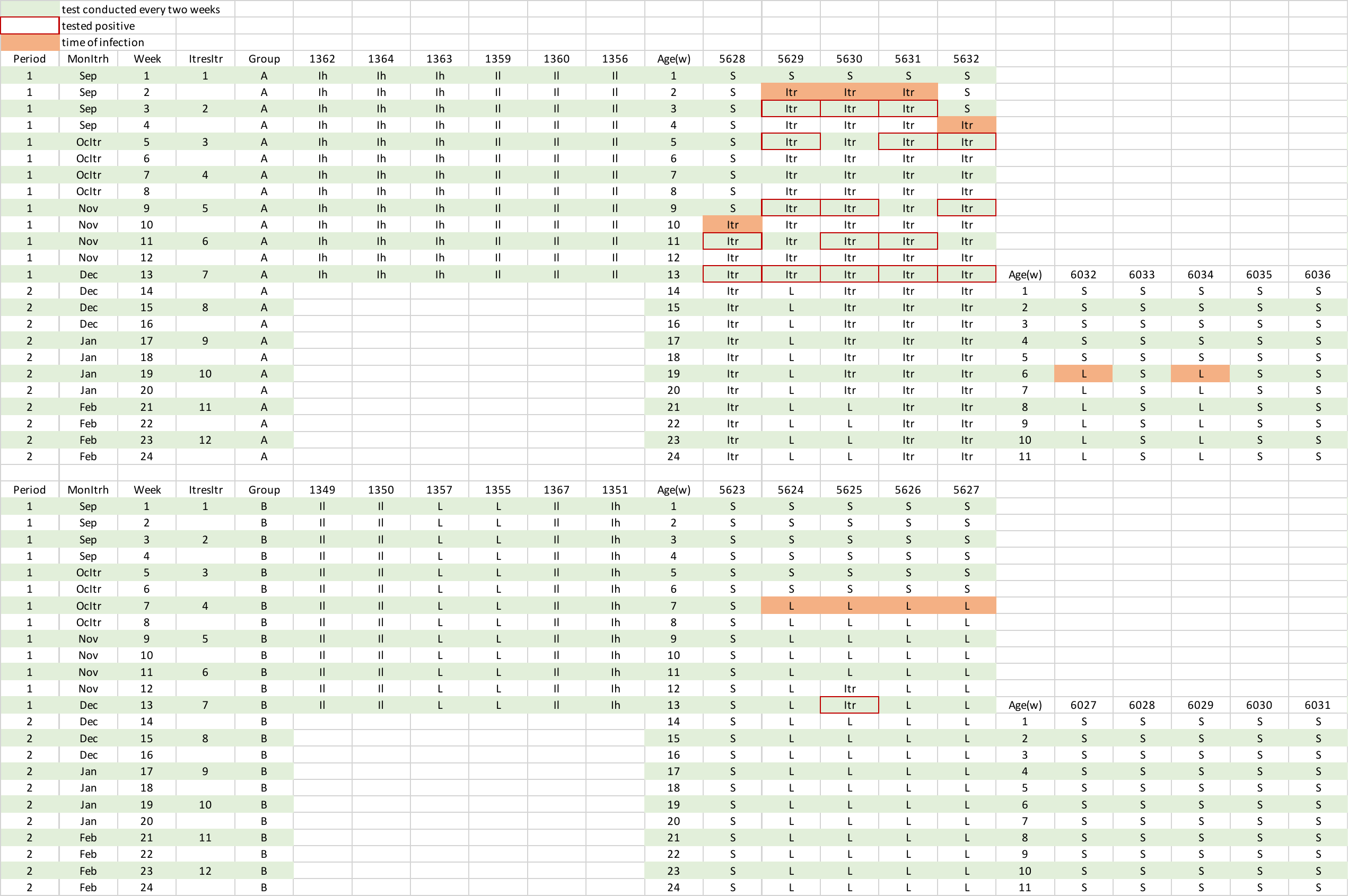
