## Supplemental materials S1-S4 files, and will be used for the link to the file on the preprint site for "Quantifying key parameters of environmental transmission and age-specific susceptibility for *Mycobacterium avium* subspecies *paratuberculosis (MAP)*": S2 File.docx

**S2 File. Detailed Schemes of Transmission Rates Based Model A and Shedding Rates Based Model B**

This supplementary file provides a detailed overview of the two models used in our study. Model A captures infectivity differences by varying the transmission rate parameters across different shedding states, whereas Model B captures these differences by varying shedding rate parameters. Both model A and B include five host compartments: susceptible individuals ($S_{a}(t)$), transient shedder ($Itr_{a}(t)$), latently infected ($L_{a}(t)$), low shedder($Il_{a}(t)$), and high shedder($Ih_{a}(t)$), all indexed by age $a$ at time $t$. The environmental compartment differs between the models: in Model A, all infectious states shed at the same rate ($sh$) but generate different types of environments ($E_{Itr}(t)$, $E_{Il}(t)$, $E_{Ih}(t)$) with varying transmission rates ($\beta_{Itr}$, $\beta_{Il}$, $\beta_{Ih}$), while in Model B, there is only one type of environment with a constant transmission rate ($\beta$), but shedding rates differ by infectious state (${sh}_{Itr}$, ${sh}_{Il}$, ${sh}_{Ih}$).

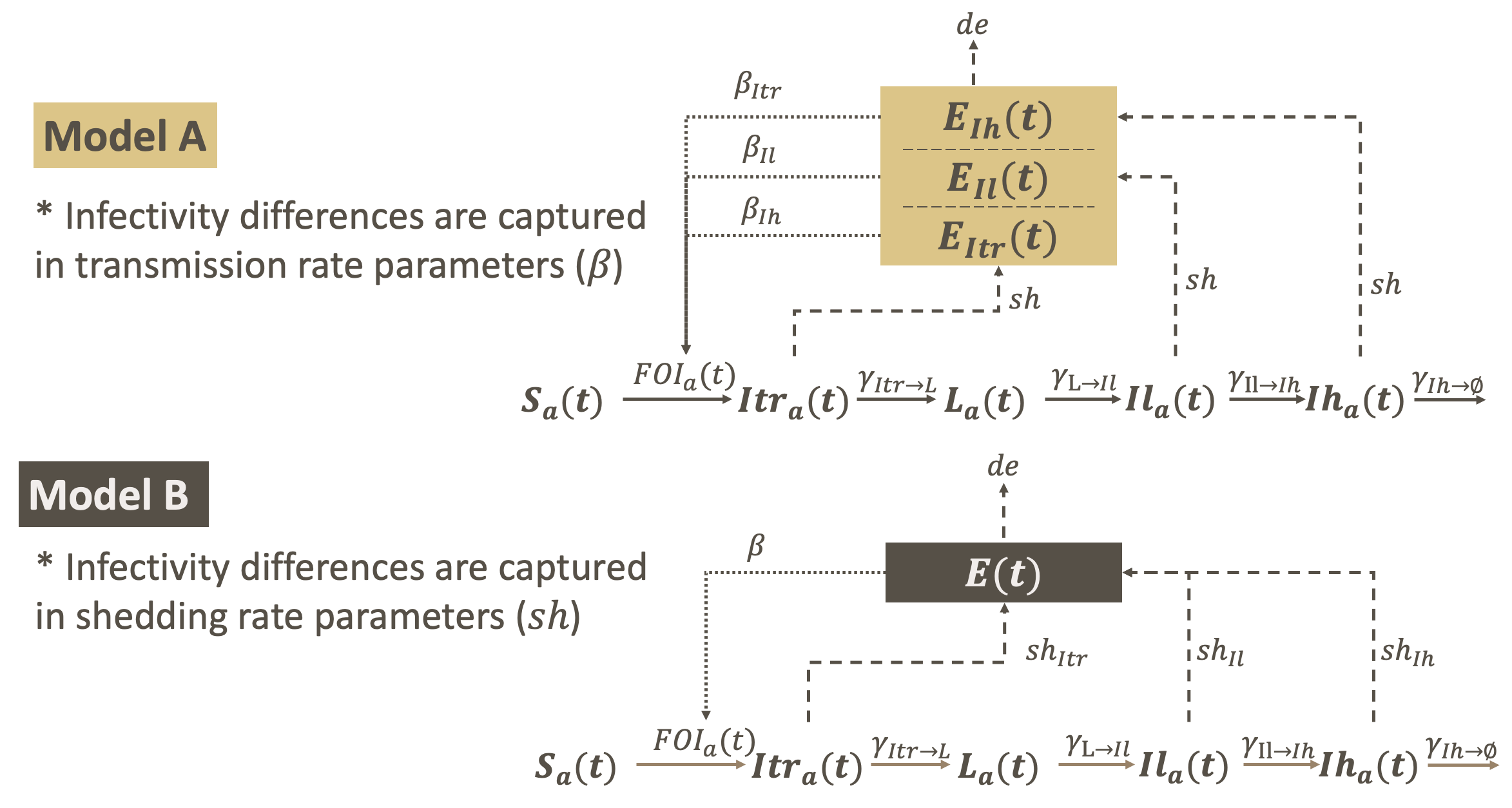

**S1 Figure. Scheme of Compartmental Model A and Model B.** $E_{state}\left( t \right)$ or $E\left( t \right)$represent environmental contamination level at time $t$ in Model A or B. $S_{a}(t)$*,* ${Itr}_{a}(t)$*,* $L_{a}(t)$*,* ${Il}_{a}(t)$*,* ${Ih}_{a}(t)$ represent number of susceptible, triansient shedder, latently infected, low shedder, high shedder aged $a$ at time $t$. ${FOI}_{a}(t)$ represent force of infection rate of a susceptible aged $a$ at time $t$, while $\gamma_{state1\to state2}$ represent transition rate from $state1$ to $state2$*. The diagram shows solid arrows for transitions between individual states, dashed arrows for *MAP* shedding and decay in the environment, and dotted arrows representing the exposure of susceptible calves.

*In the parameter estimation, individuals’ infection states were observed rather than modeled through a compartment model. The transition rates (𝛾) shown in the figure were not used in this study.
