## Supplemental materials S1-S4 files, and will be used for the link to the file on the preprint site for "Quantifying key parameters of environmental transmission and age-specific susceptibility for *Mycobacterium avium* subspecies *paratuberculosis (MAP)*": S3 File.docx

**S3 File. Validation of estimated decay rate parameter using culture-based experimental data on *MAP* viability in cattle excretions.**

To interpret and verify our estimate, we conducted a literature review focusing on *MAP* survival studies in various contaminants, primarily based on fecal culture. Since different studies used varied instructors (e.g., half-life values, decimal reduction time, expected survival time), we standardized them to conduct the comparison.

The decay constant, which is denoted as $de$ in this research, can be calculated using the exponential decay formula:

$$N\left( t \right)=N_{0}\cdot e^{-de \cdot t}$$

The half-life ($t_{1/2}$) is calculated using:

$$t_{1/2}=\frac{Log(2)}{de}$$

For some research, decimal reduction time(D-value) was given. Based on the definition, half-life value can also be calculated from D-value:

$$de=\frac{Log(10)}{D}$$

The mean survival time ($t_{surv}$) is the reciprocal of the decay constant:

$$t_{surv}=\frac{1}{de}$$

**Table S2. Culture-based experimental data on *MAP* viability in cattle excretions.**

| **Material** | **Condition** | **decay constant** | **half-life value** | **Mean survival time (wk)** | **REF** |
| --- | --- | --- | --- | --- | --- |
| Cattle slurry | 5 ℃ | 0.353 | 1.96 | 2.83 | (1) |
| Cattle slurry | 15℃ | 0.908 | 0.764 | 1.10 | (1) |
| Cattle slurry | 35 ℃ | 1.83 | 0.379 | 0.546 | (2) |
| Cattle slurry | 53-55 ℃ | 54.3 | 0.0128 | 0.0184 | (2) |
| Amitraz cattle dip fluid | 22 ℃ | 2.56 | 0.271 | 0.390 | (3) |
| urine and feces in tap water (ph 7) | 38 ℃ | 0.256 | 2.71 | 3.92 | (4) |
| urine and feces in tap water (ph 5 or 8.5) | 38 ℃ | 0.297 | 2.33 | 3.37 | (4) |
| Distilled water |  | 0.177 | 3.91 | 5.65 | (5) |
| Cattle feces exposed to unlight | 20 ℃ | 0.693 | 1.00 | 1.44 | (6) |
| Cattle feces shaded | 17.4 ℃ | 0.0898  (0.0783, 0.147) | 7.71  (4.71, 8.86) | 11.1  (6.80, 12.8) | (6) |
| Cattle feces at site Armidale | 3.6–19.7℃ | 0.118  (0.0638,0.693) | 5.86  (1.00,10.9) | 8.46  (1.44,15.7) | (6) |
| Cattle feces at site Bathurst | 6.8–19.8℃ | 0.242  (0.103,0.693) | 2.86  (1.00,6.71) | 4.13  (9.69,15.7) | (6) |
| Cattle feces at site Condobolin | 10.2–24.4℃ | 0.242  (0.0783,0.693) | 2.86  (1.00,8.86) | 4.13  (9.69,12.8) | (6) |
| Cattle feces at site Broken Hill | 11.9–24.3℃ | 0.693  (0.242,0.693) | 1.00  (1.00,2.86) | 1.44  (1.44,4.13) | (6) |
| Lab strain-Milk during the process of cheese making |  | 0.211 | 3.30 | 4.76 | (7) |
| soured milk products (yogurt, kevir etc.,) | 4 °C | 0.320 | 2.75 | 3.97 | (8) |
| fecal pellet samples collected from boxes in the 100% shade treatment | 10-22 ℃ | 0.189 | 3.66 | 5.29 | (9) |
| fecal pellet samples collected from partially shaded pasture boxes | 10-22 ℃ | 0.461 | 1.51 | 2.17 | (9) |
| fecal pellet and soil samples from partially shaded plots | 10-22 ℃ | 0.201 | 3.44 | 4.98 | (9) |

Our study estimated a *MAP* decay constant of 0.424 (95% CI: 0.0260, 2.89) per week, aligning with several previously reported values. Comparatively, Whittington et al.(9) observed decay constants of 0.461 per week for fecal pellet samples collected from partially shaded pasture boxes under 10-22 ℃. While Eppleston et al.(6) found decay constants of 0.0898-0.693 per week for Cattle feces at various sites. Additionally, Olsen et al.(2) and Jorgensen (1) reported similar decay rates of *MAP* in cattle slurry from 0.353 to 1.83 under 5-35 ℃.

Overall, our finding is well supported by the decay rates observed in general cattle farm environments, particularly in the presence of cattle feces. However, a limitation of our study is the relatively wide 95% confidence interval for the decay constant, which introduces some uncertainty in our estimate. This limitation could be addressed by fitting our model to a larger dataset from additional transmission experiments, thereby refining the parameter estimates and improving the precision of our findings.
