## Supplemental materials S1-S4 files, and will be used for the link to the file on the preprint site for "Quantifying key parameters of environmental transmission and age-specific susceptibility for *Mycobacterium avium* subspecies *paratuberculosis (MAP)*": S4 File.docx

**S4 File.** **A Comparative Analysis with A Previously Published Transmission Model**

**S4.1 Introduction**

Several transmission models have been developed to simulate the transmission processes of *Mycobacterium avium subspecies paratuberculosis* (MAP), either within experimental setups or across populations(1-11).

Marcé et al.’s transmission model is selected as a baseline for comparison due to its detailed approach, adaptability, and widespread application. In this file, we construct a dose-response environmental transmission model following Marcé’s methodology and parameter values(1). Based on constructed dose-response transmission model, we replicate the scenario study and model fit analysis.

This comparison has two primary aims: (i) to evaluate our environmental transmission model in the main text—built on a novel mathematical approach introduced in 2023 (13)—against a previously published model to gain insight into its relative performance, and (ii) to demonstrate the application and interpretation of our model with varying assumptions on transmission mechanisms.

The following results are not directly taken from the original publications; rather, we developed the model and conducted the analysis in accordance with the published methodologies and parameters in the original publications.

**S4.2 Materials & Methods**

We choose the model setup, where in a dairy farm, animals are categorized into age groups: newborn, unweaned, weaned, young heifer, bred heifer, and cow. Infected animals are further classified into three infectious states: $I_{T}$ (transiently infected), $I_{M}$ (moderately infected), and $I_{H}$ (highly infected). To calculate the total amount of *MAP* shed per week for all infected individuals within a specific age group and infectious state ($Q_{state}^{group, faeces}$), use the following equation:

$$Q_{state}^{group, faeces}\left( t \right)=\sum_{1}^{{Tot}_{state}^{group}(t)} (7\times f_{group}\times{Map}_{state}^{faeces})$$

where ${Tot}_{state}^{group}(t)$ denotes the total number of infected animals in the specific age group and infectious state at time $t$. $f_{group}$ represents the average feces production in kilograms per day, which varies by age group:

$$f_{newborn}=0.4$$

$$f_{unweaned}=0.4$$

$$f_{weaned}=4.1$$

$$f_{youngheifer}=7.5$$

$$f_{bredheifer}=22.5$$

$$f_{cow}=22.5$$

${Map}_{state}^{faeces}$ represent the *MAP* shed per kilogram of feces and is infectious state-dependent:

$${Map}_{I_{T}}^{faeces}={10}^{6}\times beta(8.8,19)$$

$${Map}_{I_{M}}^{faeces}={10}^{\left( 4+10\times beta\left( 2.65,17 \right) \right)}$$

$${Map}_{I_{H}}^{faeces}={10}^{\left( 8+10\times beta\left( 2,17 \right) \right)}$$

Here, $beta\left( a,b \right)$represents a beta-distributed random variable with parameters $a$ and $b$.

To model the amount of *MAP* present in the environment at time $t$ ($E_{state}^{group, faeces}\left( t \right)$), considering both the natural decay of *MAP* and the shedding of *MAP* from infected animals, use the following equation:

$$E_{state}^{group, faeces}\left( t \right)=E_{state}^{group, faeces}\left( t-1 \right)\times\left( 1-\mu_{indoor} \right)+Q_{state}^{group, faeces}(t)$$

where $\mu_{indoor}$ represents the decay rate of *MAP* in the indoor environment, which equals to 0.4 per week.

The relative susceptibility of a recipient calf at age $a$, which exponentially decreases with age at a rate of 0.1 per week, follows the equation:

$$relativeS\left( a \right)=e^{-0.1 a}$$

The infection probability of exposing a recipient calf of age $a$ to a contaminated environment at time $t$ ($E_{state}^{group, faeces}\left( t \right)$) was calculated using the following formula:

$$Infection probability\left( a,t \right)=1-e^{\left( relativeS\left( a \right){\frac{\beta E_{state}^{group, faeces}(t)}{N\left( t \right)}}/\alpha\right)}$$

where $\beta$ represents the transmission rate parameter for the indoor environment per week, which equals to $5.0\times{10}^{-5}\times7$. $N(t)$ represents the total number of animals present in the environment at time $t$. $\alpha$ represents the infectious dose, which equals to $1\times{10}^{6}$.

**S4.3 Results**

The scenario study, replicating the four main objectives from the text, showed that the total *MAP* concentration in the environment increased rapidly before stabilizing. However, there was a significant difference between high shedders (BredheiferH and CowH) and other infectious states. Specifically, only high shedders had a non-zero probability to infect a susceptible individual during one week (others being zero to two decimal places, i.e. probability <0.005) across all scenarios. Although differences are observed between M shedders and T shedders (as shown in the third column of Figure S1, excluding H shedders), these differences are not practically significant due to the very low infection probabilities.

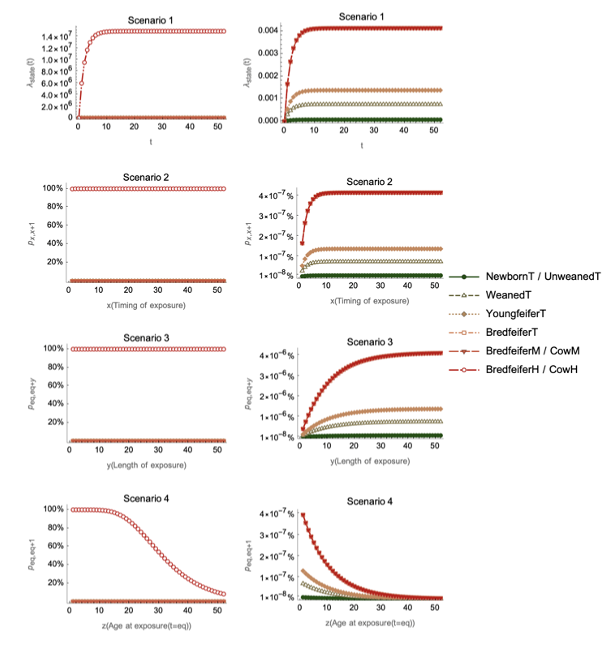

**Figure S2. Scenario Study Results of Dose-Response Transmission Model.** In compare with the results shown in main text Figure 5&6, the left column presents the overall outcomes from the scenario study using the dose-response transmission model. Since all results, except for BredheiferH/CowH, overlap at 0, data excluding BredheiferH/CowH are shown in the right column to highlight differences among individuals in various states and groups. Note that the y-axis ranges differ between the first two columns and the third column.

Additionally, we performed a model fit analysis to compare the predicted infection probabilities with the observed infections from the original experiments. The predicted number of new cases over the entire period of experiments A1, A2, B1, and B2 were 5.00, 0.00, 3.98, and 0.00, respectively, compared to the observed values of 5, 2, 4, and 0. Furthermore, the predicted infection probability was calculated and compared. As illustrated in Figure S3, the models fit well for Experiments A1 and B1. However, in Experiment B2, the model predicted no new infections, while 2 out of 5 recipients were actually infected, suggesting that calf-to-calf transmission may be underestimated by the model.

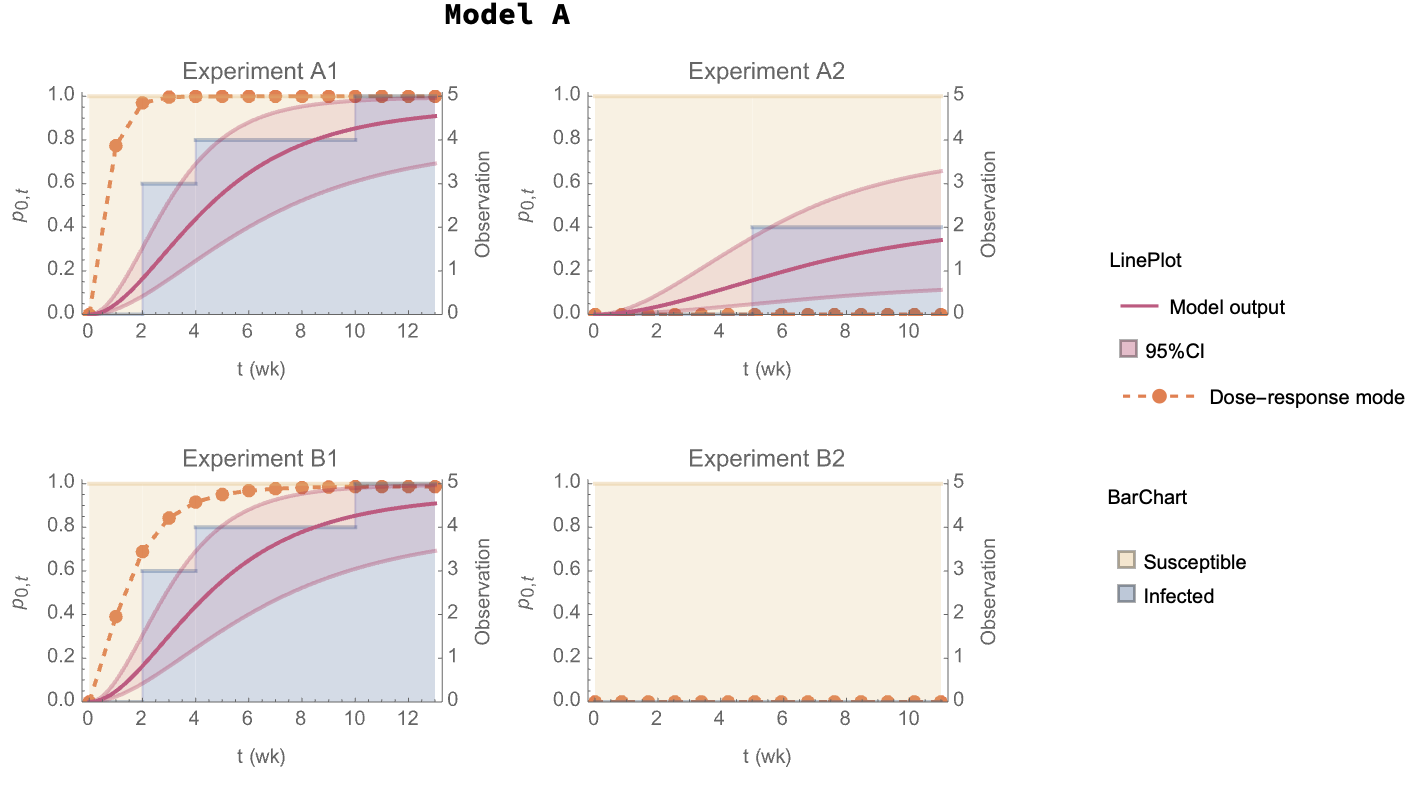

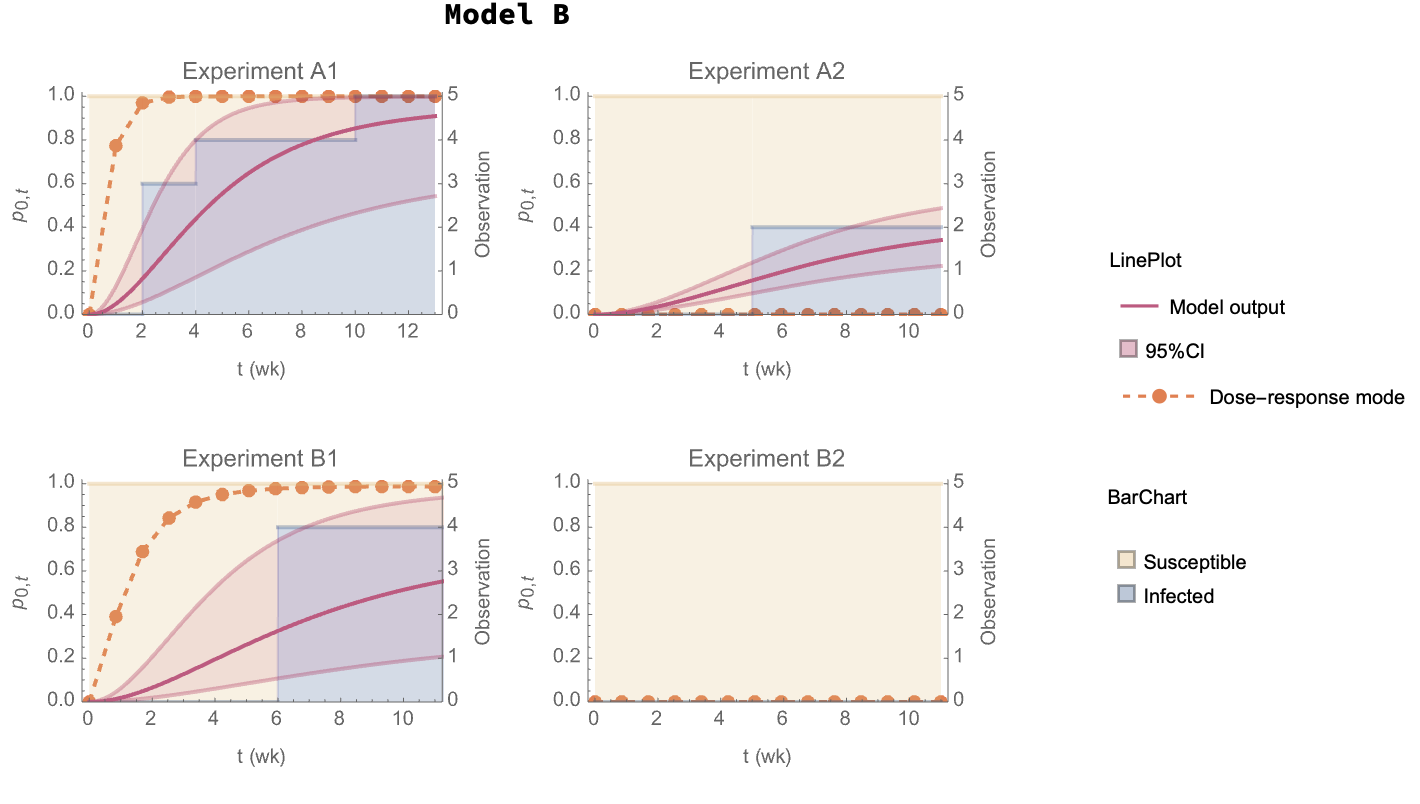

**Figure S3. Comparison of Predicted Infection Probability with Experimental Observations.** The solid curve and shaded area represent the predicted infection probability from (0,t) and their 95% confidence interval from model present in main text, while the dotted lines represent prediction from dose-response transmission model in S4 file. The x-axis denotes time in weeks, with the left y-axis showing infection probability and the right y-axis showing the number of individuals.

**S4.4 Discussion**

This file compares the estimated parameters and environmental transmission models developed in main text to a dose-response transmission model commonly used in previous studies. Key parameters compared include age-specific susceptibility rates, transmission rates, and *MAP* decay rate.

The dose-response model categorizes animals by age (newborn to cow) and by infectious states: transiently infectious ($I_{T}$), moderately infectious ($I_{M}$), and highly infectious ($I_{H}$). The total *MAP* shed by infectious animals is calculated based on feaces production, which varies by age group, and the amount of *MAP* shed per kilogram of feaces, which depends on the infectious state.

The model incorporates natural *MAP* decay in the environment and uses a age-specific susceptibility function to calculate the infection probability of calves exposed to contaminated environments. The scenario study showed that high shedders significantly increased environmental *MAP* contamination and infection probabilities. However, there are significant differences in contributions from moderately shedders and transiently infectious shedders. This observation is further supported by the model fit analysis, where the predicted infection probabilities for cow-to-calf transmission generally aligned with experimental data. However, in calf-to-calf transmission (Experiment B2), the dose-response model underestimated the probability. For instance, with five recipients in each experiment, the number of new infections predicted by our environmental transmission model was 4.81, 2.12, 3.58, and 0.00, while the dose-response model estimated 5.00, 0.00, 3.98, and 0.00 new infections, respectively. In comparison, the actual observed infections were 5, 2, 4, and 0.

By comparing the environmental transmission model (in main text) and dose-response model( in S4 file), the study highlights the strengths and limitations of each approach, providing a case study on how models using different mechanisms can be compared with each other and evaluated against experimental data.
